# Protein nanobarcodes enable single-step multiplexed fluorescence imaging

**DOI:** 10.1101/2022.06.03.494744

**Authors:** Daniëlle de Jong-Bolm, Mohsen Sadeghi, Guobin Bao, Gabriele Klaehn, Merle Hoff, Lucas Mittelmeier, F. Buket Basmanav, Felipe Opazo, Frank Noé, Silvio O. Rizzoli

## Abstract

Multiplexed cellular imaging typically relies on the sequential application of detection probes, such as antibodies or DNA barcodes, which is complex and time-consuming. To address this, we developed here protein nanobarcodes, composed of combinations of epitopes recognized by specific sets of nanobodies. The nanobarcodes are read in a single imaging step, relying on nanobodies conjugated to distinct fluorophores, which enables a precise analysis of large numbers of protein combinations.

## Main text

Fluorescence imaging is one of the most powerful tools for cellular investigations, but its potential to reveal multiple targets has been rarely fulfilled, due to difficulties in labeling many molecules simultaneously, or in separating multiple fluorophores spectrally ^1^. One potential solution has been the introduction of multiplexing by sequential labeling, in which reagents carrying the same fluorophore are added and removed sequentially. This can be achieved by fluorophore bleaching (for example in toponome mapping ^2^), by antibody removal using harsh buffers, or by probe removal by extensive wash-offs (for example maS^3^TORM ^3^ or DNA-PAINT ^4^). While these approaches have been used to investigate samples from cancer cells to synapses, they involve long-lasting and challenging experiments, implying that a simpler solution is desirable.

To approach this problem, we started from the idea that every microscope has a handful (*n*) of spectrally distinguishable channels, with which *n* specific labels should be differentiated relatively easily. The number of possible combinations of labels is substantially higher than *n*, since each label can be present or absent (“on/off” signals), which leads, in theory, to 2^*n*^ combinations, as in a conventional barcode. As the “all labels absent” combination is useless for practical purposes, the actual number of targets that could be differentiated becomes 2^*n*^-1. Therefore, this barcoding approach could be used to strongly enhance the number of targets that can be analyzed simultaneously using a limited number of channels. So far, it has been used for cell identification by fluorescence cell sorting (FACS ^5^), using antibody detection, but could not yet introduced in the domain of conventional microscopy. Imaging the different label combinations using antibodies is almost impossible, due to problems with steric hindrance caused by the large antibody size, label clustering induced by the dual binding capacity of the antibodies, or limited epitope availability due to poor penetration into the cells ^6,7^.

To solve this issue, we relied on epitope recognition by nanobodies (single-domain camelid antibodies), which are monovalent and substantially smaller than antibodies ^8,9^. To establish a proof-of-principle approach, we engineered proteins that contain a combination of five genetically-encoded epitopes that are recognizable by nanobodies. These combinations were termed “nanobarcodes”, and consisted of the following epitopes. First, a reference epitope was added to all our nanobarcodes, in the form of the ALFA-tag ^10^, enabling us to detect every nanobarcode, irrespective of what other epitopes are present. The other four epitopes were present only in subsets of all nanobarcodes: mCherry(Y71L) and GFP(Y66L), both mutated to generate non-fluorescent variants ^11^ and two different short sequences found at the C-terminus of human α-synuclein^12^ (termed here syn87 and syn2). These four epitopes were engineered, in different combinations, into the sequences of different proteins, and were then revealed using the respective fluorescently-labeled nanobodies (NbRFP, NbEGFP, NbSyn87, NbSyn2). As designed, all the epitopes were easily detected in immunocytochemistry (Fig. 1).

**Figure 1:**
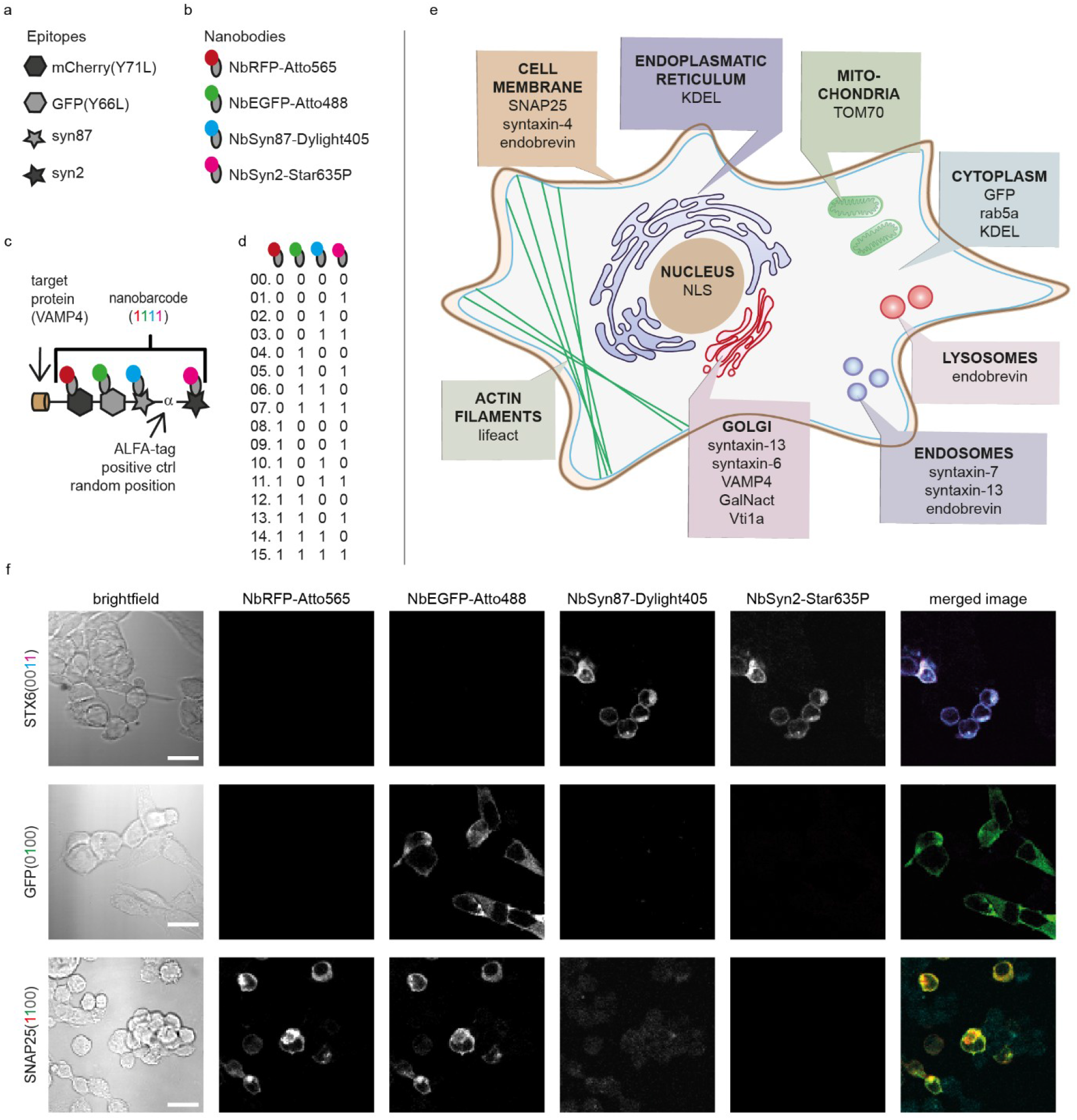
Design of protein constructs with nanobarcodes using four nanobody epitopes. **a, b:** Scheme of the four nanobarcode epitopes (a) and the fluorescent nanobodies used for recognizing them (b): NbSyn87-Dylight405 in cyan, NbGFP-Atto488 in green, NbRFP-Atto565 in red, NbSyn2-Star635P in magenta. **c:** Design of the protein construct VAMP4(1111). Each protein construct contains a target protein (the protein to identify) and a nanobarcode. In this example, the target protein is VAMP4 and its nanobarcode contains the following non-fluorescent epitopes: mCherry (Y71L), GFP (Y66L), syn87, and syn2. The ALFA-tag ^10^ is present for testing purposes. See Extended Data 1 for further sequence information. Nanobarcode epitopes recognized by fluorescent nanobodies are shown as “ones” in pseudo-colors that correspond to the fluorophores used. **d:** Nanobarcodes, fifteen in total, resulting from a binary combination of four nanobarcode-epitopes. Epitopes from left to right: mCherry(Y71L), GFP(Y66L), syn87 and syn2. The nanobody scheme is the same as in b. **e:** The expected cellular protein distribution for the proteins used, according to the literature. **f:** Nanobarcode-based identification of the proteins STX6(0011), GFP(0100) and SNAP25(1100). The pseudo-colors for merged images correspond to the fluorescence channels of the nanobodies: NbRFP-Atto565 in red, NbGFP-Atto488 in green, NbSyn87-Dylight405 in cyan and NbSyn2-Star635P in magenta. Scale bar: 20 µm.

We implemented the nanobarcodes in 15 different proteins (2^4^-1), according to the schemes shown in Fig. 1a-d. We targeted proteins mostly from the secretory pathway, such as Vesicle-Associated Membrane Proteins (VAMPs) and Syntaxins (for a schematic topology of all protein constructs, see Extended Data Fig. 1), as indicated in Fig. 1e. To validate our approach, we expressed the barcoded proteins and compared their nanobarcode images to simple immunostainings for the respective endogenous protein epitopes (Extended Data Fig. 2). We then verified the accuracy of the nanobody-based staining (Extended Data Fig. 3-4). The approach appeared to function well, as illustrated by the images in Fig. 1f, in which the nanobarcodes can be easily differentiated by the human observer.

To facilitate the analysis of the nanobarcode images, we developed an artificial neural network to connect the combined fluorescence output of the nanobarcode sequences to the identity of the respective labelled proteins (Fig. 2a, Extended Data Fig. 5). We formulated the analysis as a pixel-wise classification problem, for which the deep learning strategy is especially well-suited. After meticulous training of the network (see Methods) we analyzed its performance based on (i) hold-out test sets, and (ii) full images of single-transfected samples with known barcoded proteins. The latter analysis resulted in a relatively high accuracy, considering the strong criterion of pixel-wise true identification (Fig. 2b-d).

**Figure 2:**
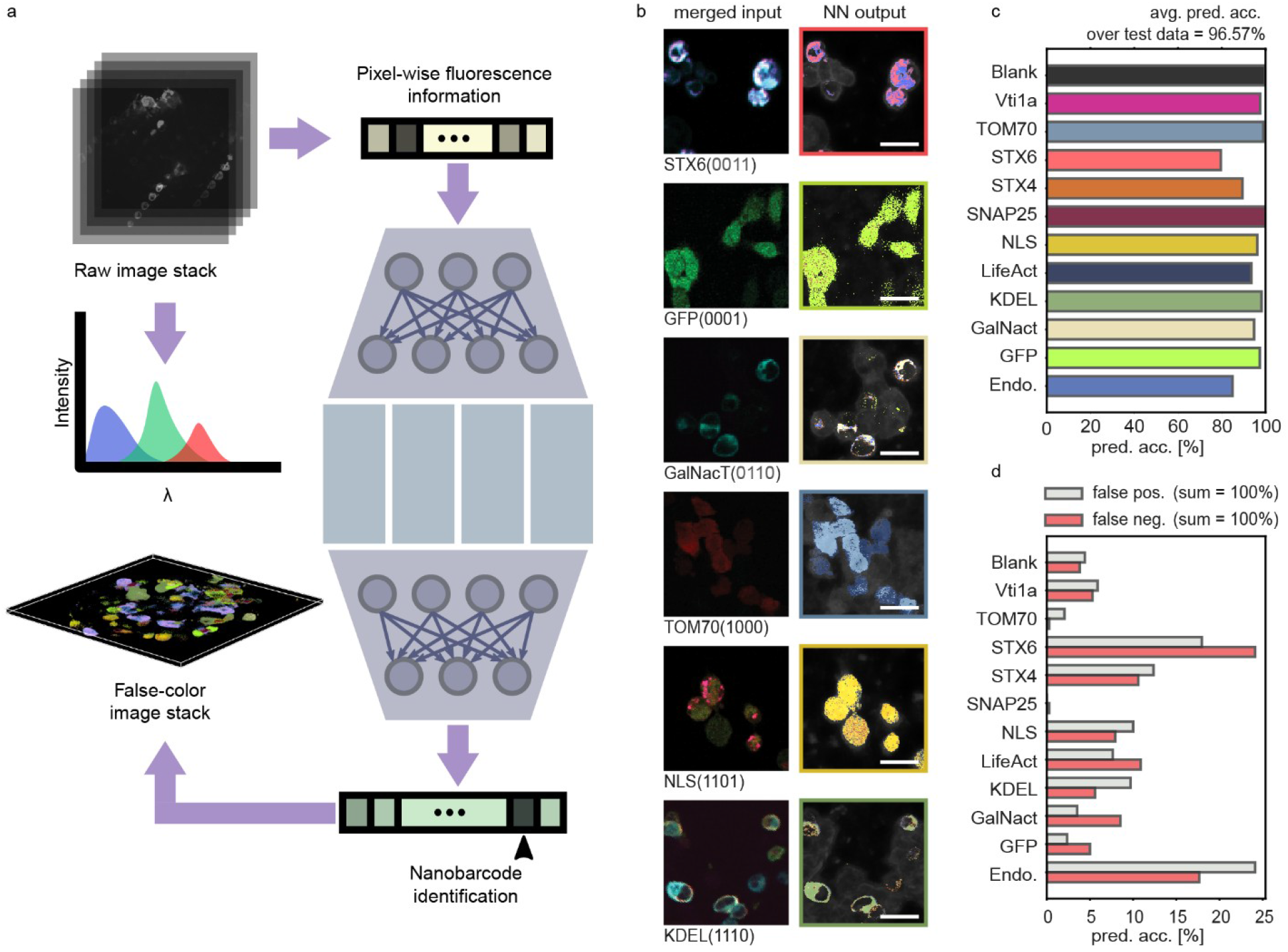
Neural-network-based identification of nanobarcode proteins. **a:** Schematic of the neural network used for identification of nanobarcodes from pixel-wise fluorescence information. Brightness values across all emission channels are fed to the network as input, which in turn has been trained to predict the probability of this information pertaining to a specific nanobarcode, or a blank pixel. The trained network can readily be applied to full micrographs as well as stacks of images to produce false-color outputs illustrating spatial distribution of proteins (further details in Extended Data Figure 5). **b:** Example images of HEK293 cells transfected with specific nanobarcodes. To account for all possible emission features (including bleed-through), we acquired eleven frames for each area, consisting of the following: 405nm excitation, with emission windows in blue, green, red, deep red; 488 nm excitation, with emission windows in green, red, deep red; 561 nm excitation, with emission windows in red and deep red; 633 nm excitation, with an emission window in deep red; brightfield. The panels in the left column show an overlay of the four brightest frames: 405 nm excitation, blue emission (in cyan); 488 nm excitation, green emission (in green); 561 nm excitation, red emission (in red); 633 nm excitation, deep red emission (in magenta). False color neural network output images are shown in the right column of a. **c:** Prediction accuracy of the neural network over a hold-out test dataset. **d:** False positive and false negative protein identifications (as percentage of all false predictions). For further details about the experimental procedures and neural network analysis, see the Methods section. For practical implementation purposes, we concentrated here on a subset of the labeled proteins, which were also used for the Nrxn/Nlgn experiments in Fig. 3. Scale bars: 20 µm.

After ensuring that the network could identify proteins with satisfactory precision, we were ready to apply it to samples with withheld combinations of transfected cells. To apply this analysis to a relevant biological problem, we turned to the analysis of a set of cell adhesion molecules that are essential in neuronal cell biology: neurexins (Nrxns1-3) and neuroligins (Nlgns1-4). Heterogeneous binding of Nrxns to Nlgns is crucial for synapse formation, and the absence of functional Nrxns and/or Nlgns variants is lethal ^13,14^. Binding between Nrxns and Nlgns is dependent on the alternative splicing of these molecules, resulting in a complex pattern of interactions ^15^. Nrxn/Nlgn binding is subject to detailed and poorly known regulation, with the interaction of specific partners being affected by neuronal plasticity and by local conditions. Aberrant interactions are also known to form, most prominently in the case of autaptic cultures consisting of isolated neurons that form synapses onto their own dendrites. Interactions between the two sets of molecules are typically investigated by introducing single splicing variants into cells, followed by a one-by-one comparison of binding properties and/or interactions between Nrxn/Nlgn pairs ^16, 17,18,19,20^. This type of analysis can pinpoint the interactions with the highest affinity, but they do not necessarily recapitulate the *in vivo* situation, in which cells expressing specific molecules navigate a multi-cellular tissue containing numerous potential partners, before settling on a specific interaction. Ideally, cells carrying different Nrxns and Nlgns should be exposed to each other simultaneously, in a multi-cell competition, to enable individual cells to test different potential partners, as in living tissues.

We therefore applied the nanobarcoding tools to this problem (Extended Data Fig. 6 and 7). We co-expressed different Nrxns and Nlgns with specific nanobarcoded proteins (Extended Data Fig. 7), and we then developed a cell-seeding assay that allows us to map all of the respective Nrxn/Nlgn interactions (Fig. 3). We applied this assay to four β-Nrxns and seven Nlgn isoforms: Nrxn-1ß (SS#4(+)), Nrxn-1 (SS#4(-)), Nrxn-2ß (SS#4(-)), Nrxn-3ß (SS#4(-)), Nlgn1(-), Nlgn1 (SS#B), Nlgn1 (SS#AB), Nlgn2 (-), Nlgn2 (SS#A), Nlgn3(WT), Nlgn4 (WT) (see also Extended Data Info.)

**Figure 3:**
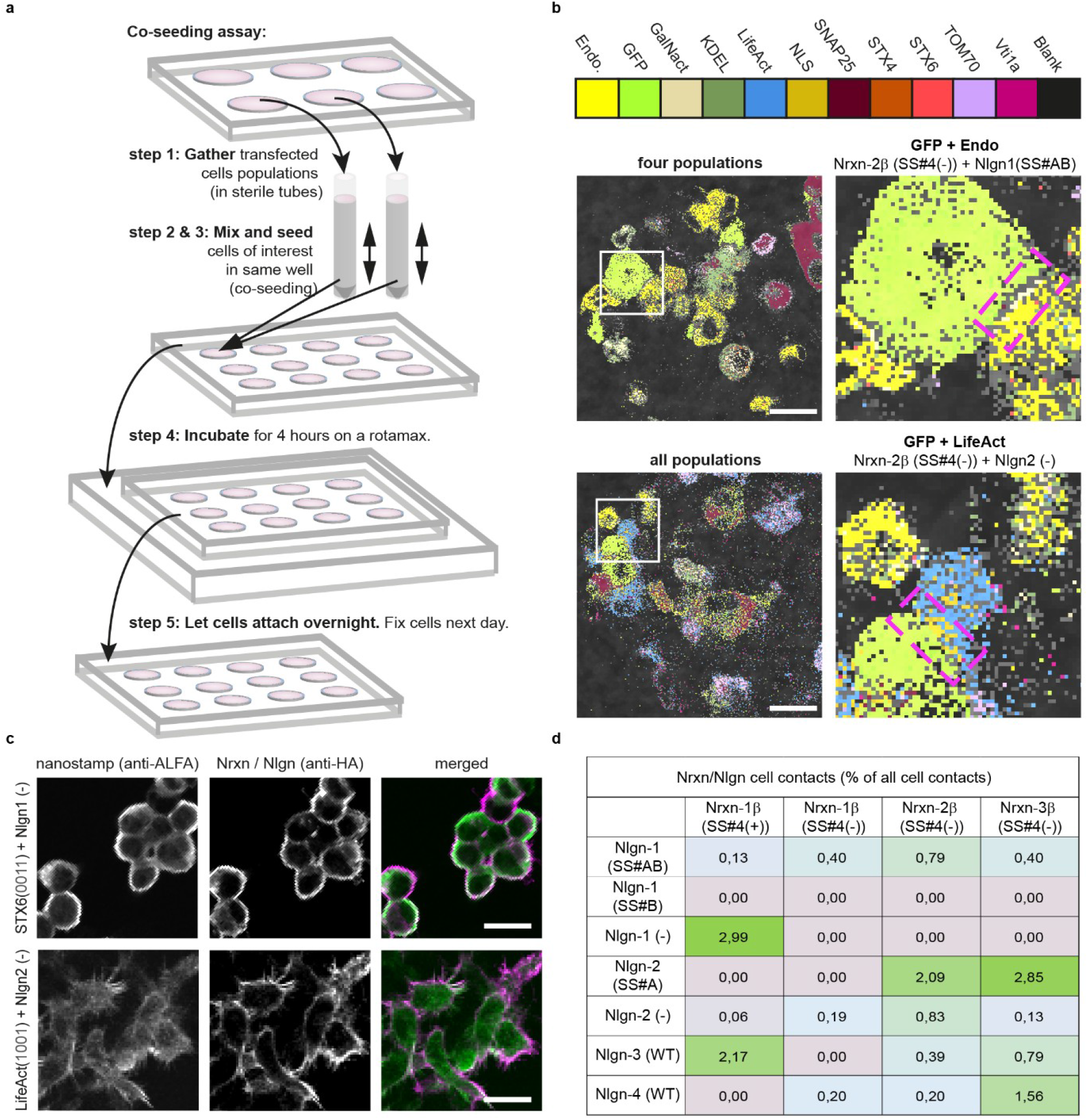
Multiplex identification of proteins using a neural-network-based spectral analysis. **a:** Experimental design of a co-seeding assay including 11 different cell types, labeled with specific nanobarcodes (see Methods section for details). **b:** Example of a Nrxn-2ß (SS#4(+))/Nlgn-1 (SS#AB) and a Nrxn-2ß (SS#4(-))/Nlgn-2 (-) pair. **c:** Overlay of cells containing nanobarcode proteins and Nrxn or Nlgn positive cells. Nanobarcode-proteins are shown in green (anti-ALFA-Atto488). Nrxn or Nlgn isoforms are shown in magenta (anti-HA & anti-goat-Cy3). See Extended Data Fig. 6 for example images of all proteins. Scale bars: 20 µm. **d:** Interaction preferences of Nrxn/Nlgn isoforms. 4569 cell contacts, 147 images, 4 independent co-seeding experiments. The Nrxn/Nlgn codes, such as SS#4(+) refer to the respective splicing sites of the proteins, according to the literature.

From the total number of cell contacts made by each Nrxn- or Nlgn-positive cell, we calculated the percentage of specific Nrxn/Nlgn pairs (Fig. 3b-d, Extended Data Info.). We found that some specific combinations are substantially more likely than others (Fig. 3d). For example, we regularly identified ß-Nrxn1^ss4+^/Nlgn3^WT^ pairs, which is surprising, since Nlgn3 is thought to have a lower affinity for Nrxns than the Nlgn1 and Nlgn2 isoforms ^20^. In addition, three other Nrxn/Nlgn pairs were observed regularly: Nrxn1ß (SS#4(+)) /Nlgn1(-), Nrxn2ß (SS#4(-)) /Nlgn2 (SS#A) and Nrxn3ß (SS#4(-))/Nlgn2 (SS#A), which are compatible with the previous literature, albeit none are known to be of particularly high affinity. This implies that such an assay should be used for testing further the Nrxn/Nlgn interactions, especially as it is able to take into account not only the molecular binding, but also the further dynamics that are induced by binding, such as molecular endocytosis and trafficking ^21^

We conclude that the nanobarcoding technology is feasible in conventional microscopy assays. However, we did not exploit this assay in super-resolution, although it should be suitable for this approach, especially as the probes used (nanobodies) have been heavily used in super-resolution for a decade (e.g. ^22^). A potential limitation is the use of genetic encoding, but the current developments in CRISPR/Cas technologies should render this approach not overly difficult. In addition, the sequences (barcodes) could be expressed, purified and linked to secondary nanobodies, which are applied to reveal primary antibodies in immunocytochemistry and are inherently multiplex-able ^23^, thereby extending the assay to many protein targets. Moreover, since many other barcode epitopes could be used, our approach should have a large application range in the field of cellular biology and proteomics.

## Supporting information

Methods and Protocols

## Acknowledgments

This project receives funding from the European’s Union Horizon 2020 Horizon research and innovation programme under grant agreement No 964016 (FET-OPEN Call 2020, IMAGEOMICS project). The authors acknowledge Ms. Christina Zeising for helping with general laboratoray procedures. M.S. and F.N. received financial support from Deutsche Forschungsgemeinschaft (DFG) through grants CRC 958/Project A04 and CRC 1114/Project C03. F.N. was additionally supported by European Research Commission grant ERC CoG 772230 and the Berlin Institute for Foundations in Learning and Data (BIFOLD). F.B.B. was supported by the Deutsche Forschungsgemeinschaft (DFG) through Cluster of Excellence Nanoscale Microscopy and Molecular Physiology of the Brain (CNMPB) and by the Campus Laboratory for Advanced Imaging, Microscopy and Spectroscopy (AIMS).

## Data Availability Statement

The data presented in this study are available on request to the corresponding author.

