## Supplementary material for "Protein nanobarcodes enable single-step multiplexed fluorescence imaging": Methods and Protocols

### Methods & Protocols

#### *In silico* design of nanobarcode proteins for protein identification

Nanobarcode proteins were designed *in silico* and consist of three main components: 1) the protein sequence, 2) up to four genetically-encoded epitopes that form the nanobarcode, and 3) the ALFA-tag [1] for testing purposes. Short flexible linkers (five amino acids long) were added in between epitopes to ensure epitope availability. We used abbreviations for the nanobarcode-proteins to ensure readability (Fig. 1B). For example, the abbreviation NLS(1101) was used for the nanobarcode-protein NLS_L2_mCherry(Y71L)_L3_GFP(Y66L)_L4_α_L5_syn2. It contains three of the four nanobarcode-epitopes, mCherry(Y71L), GFP(Y66L) and syn2, plus the ALFA-tag for testing purposes. Accordingly, the NLS construct contains four flexible linkers (L2-L5). The position of the ALFA-tag and the positions of the flexible linkers varied among the palette of proteins used, according to the characteristics of each protein. For full length sequences see Extended Info. The company GenScript® Biotech generated the pcDNA3.1(+) mammalian expression vectors containing the nanobarcode-sequences DNA, using the NheI/XhoI cloning sites.

#### Cell culture experiments with Human Embryonic Kidney 293 cells (HEK293 cells)

*HEK293 cell cultures*

HEK293 cells were cultured in a $CO_2$ incubator (37°C and 5 % $CO_2$). Cells were maintained in Dulbecco’s-Modified-Eagle’s-Medium (SIGMA-ALDRICH®, 10 % Fetal Bovine Serum (SIGMA-ALDRICH®), 2 % L-Glutamine (Gibco), 0,6 % Penicillin/Streptomycin (Lonza). When confluent, cells were briefly washed with DPBS (Dulbecco’s Phosphate Buffered Saline) and were detached from the culture dishes using 0.05% Trypsin-EDTA (1X, Gibco), before culturing them in fresh medium. Approximately 24 hours before plasmid transfection, HEK293 cells were seeded on 12-well plates containing poly-L-lysine coated glass coverslips. An exception was made for the co-seeding assay (see below), in which the cells were first mixed in solution, before finally being seeded on glass coverslips.

*Lipofectamine-based transfection of HEK293 cells*

HEK293 cells were transfected with 1 – 2 µg of plasmids per well, after mixing with 1.5 – 4 µl of Lipofectamine® 2000 (for DNA sequences, see Suppl. Info.). The optimal time window for transfection was defined based on the protein performance in the neural network analysis (see below for details about the identification procedure, Extended Data Figure 6). Tested time windows were overnight, 24 h, 48 h and 72 h.

*Co-seeding of cell suspensions containing distinct HEK293 cell populations*

For co-seeding, transfected HEK293 cells were trypsinized, washed and brought to suspension in complete medium without antibiotics. Subsequently, cells transfected with different constructs were mixed. Up to eleven different transfected cell populations were co-seeded into a single 12-well plate (CellStar®) containing poly-L-Lysine-coated coverslips. Plates with co-seeded cells were gently shaken in a humidified incubator for 1 h and then stopped, allowing cells to attach to the coverslips overnight.

**Nanobody production and coupling**

Nanobodies were custom produced (NbSyn87 and NbSyn2) or simply purchased as catalog products from NanoTag Biotechnologies GmbH (Göttingen, Germany), as described below.

**Immunocytochemistry procedure**

*Immunocytochemistry with Antibodies and or Nanobodies*

Transfected HEK293 cells were fixed with 4% PFA for 45 minutes at room temperature, followed by a short rinse in PBS and aldehyde quenching with 100 mM $NH_4Cl$ and 100 mM glycine in PBS, for 30 minutes at room temperature. Cells were permeabilized and blocked using PBS supplemented with 0.01% Triton X-100 and 2% bovine serum albumin (BSA) for 30 minutes at room temperature. Permeabilized cells were immunostained with the following fluorescent nanobodies: NbSyn87 (conjugated to DyLight 405), NbEGFP (FluoTag®-Q anti-GFP Atto488, Cat#N0301-At488-L), NbRFP (conjugated to

Atto565, sold as FluoTag®-Q anti-RFP, Cat#N0401-AT565-L), NbSyn2 (conjugated to Star635P). All nanobodies have been characterized and used in early studies (the NbEGFP and NbRFP [2] and the NbSyn2 and NbSyn87 [3–6]). Nanobodies were incubated for 1 h at room temperature in the permeabilization/blocking buffer indicated above, at final concentrations of ~70 nanogram/µl (NbEGFP), 70 nanogram/µl (NbRFP), ~70 nanomolar (NbSyn87), ~70 nanomolar (NbSyn2). Excess nanobody was thoroughly washed with PBS and coverslips were mounted on microscope slides using Mowiol. For antibody stainings, procedures were very similar but now stainings were achieved by 1 h incubation with primary antibodies followed by an 30 min. incubation with secondary antibodies (For details regarding antibodies and used concentrations, see Extended Data Table 1).

**Imaging and Image processing**

Multi-channel images were obtained using a ZEISS LSM 710 AxioObserver equipped with a ZEISS Plan Apochromat 63x oil DIC objective lens (NA 1.40). Images were acquired using 512x512 pixels, at 440 nm pixel sizes. Samples were illuminated with the following lasers: λ = 405 nm, λ = 488 nm, λ = 561 and λ = 633 nm. Fluorescence was collated using the corresponding emission filters 417-485 nm (CH1), 495-553 nm (CH2), 573-631 nm (CH3), 641-729 nm (CH4).

The following combinations of lasers (excitation) and emission channels were used for our 10-channel recordings:

- 405 nm plus CH1-4 (images 1-4),
- 488 nm plus CH2-4 (images 6-8)
- Brightfield image for validation purposes (image 5)
- 561 nm plus CH3-4 (images 9 and 10)
- 641 nm plus CH4 (image 11)

Validation images of the fluorescent protein constructs stained with primary and secondary antibodies (see Extended Data Fig. 3 and Extended Data Table 1) were obtained using an Olympus IX71 microscope equipped with an Olympus UPlanSApo 60x oil objective (1.35 NA), except for Rab5a images. These were obtained using the ZEISS LSM710 AxioObserver described above.

**Deep neural network-based protein identification**

We have designed and trained a deep neural network for the identification of proteins from confocal images (Extended Data Fig. 5). We have taken an approach similar to image segmentation, and have assigned to each pixel of the image probabilities corresponding to the presence of each of nanobarcode-proteins. In essence, the network has to learn a mapping between channel intensity values to a so-called Multinoulli probability distribution. The components of the vector $x=(x_1,\dots,x_n)^T$ represent intensities of each of the $n$ imaging channels. This vector is fed to the network as the input, producing $m$ outputs $y_1,\dots,y_m$. These values are to represent the conditional probability $P(1_i|x;\theta)$, with $1_i$ denoting the pixel containing the $i$-th nanobarcode, and $\theta$ being the parameters of the network. In order for the output of the network to be a normalized probability distribution, the last layer applies a softmax function (Extended Data Fig. 5b),

$$P(1_i|x;\theta)=softmax(y_i)=\frac{\exp(y_i)}{\sum_j \exp(y_j)}$$

The loss function, minimizing which with respect to $\theta$ constitutes the training procedure, is the negative log-likelihood of the Multinoulli probability distribution,

$$L=-\log(P(1_i|x;\theta))=-y_i+\log\left(\sum \exp(y_j)\right)$$

Maximizing the log-likelihood over the probabilities is equivalent to minimizing the cross-entropy between the data distribution and the distribution modeled by the network [[7]].

We have designed the feed forward network by stacking residual blocks (Extended Data Fig. 5b). Using residual learning allows the training of significantly deeper networks [[8]]. In addition, by increasing cardinality, i.e. inclusion of multiple parallel paths through the network, we allow for high representation power with less network depth, thus preventing vanishing gradients during the training procedure [9] (Extended Data Fig. 5b). Dense layers of the network apply an affine transformation to their input, followed by the nonlinear activation function $g$. Thus, for vectors $z$ being transformed by the network, $z_{n+1} = g\left(W_n z_n + b_n\right)$. We have used Rectified Linear Unit (ReLU) as the activation function $g$ throughout the network, and have employed the batch normalization algorithm for enhanced convergence [10] (Extended Data Fig. 5).

The data for supervised training of the network is prepared by taking micrographs including known nanobarcode-proteins. We use the $l_1$-norm of channel intensities as a measure of the total brightness of a pixel and select pixels with most reliable signals via dynamic brightness thresholding. The procedure consists of adapting a threshold based on cumulative histogram versus decreasing brightness values. The background is filtered out by automatically recognizing a sharp rise in the number of pixels. The procedure consists of adapting a threshold until a preset number of samples have been gathered. To each samples containing proteins, we gather 10 blank samples to capture the background noise in the absence of tagged proteins. For the training to cover situations where the actual signal-to-noise ratio is lower than the gathered samples, we apply a contrast augmentation to the training data. We scale the values of channel intensities in the range 0.5 to 1.5 in a stochastic manner each time the network accesses the training data. The approach results in a more robust prediction as well as less chance of over-fitting.

The network is trained via gradient descent using the AdamW algorithm [11,12]. Gradient of the loss function with respect to trainable parameters is calculated in the forward pass and the optimizer algorithm updates the parameters via backpropagation [13]. The input data are split 80%-20% into training and hold-out test datasets, with the network being trained only on the former. We monitor the training procedure by comparing the loss between training and test datasets. We stop the training loop when the test loss plateaus, which implies the network might begin to over fit to the training data.

After the training of the network is complete, full-sized images are fed to the network in a pixel-by-pixel scan. We have produced output images by assigning false colors to each protein, and using output probabilities to compose a weighted color sum per pixel (Extended Data Fig 5c). In order to account for slight variations in imaging conditions, we additionally applied trainable shifts and scales to the input brightness values. These transformations are separately trained per image in an unsupervised manner via minimizing the total entropy of the output signal. Apart from this, we have applied no other pre- or post-processing to the images.

Apart from trainable parameters, the network design contains a set of so-called hyperparameters, which in our case are the maximum layer width in each branch, number of branches, depth of each branch, and the learning rate. We have used the Adaptive Experimentation Platform (AX) to optimize hyperparameters based on the network performance, i.e. its prediction accuracy on the test dataset, via Bayesian optimization [14].

14. Ax · Adaptive Experimentation Platform. Available at: https://github.com/facebook/Ax.
