## Extended Data Figures and Tables for "Protein nanobarcodes enable single-step multiplexed fluorescence imaging"

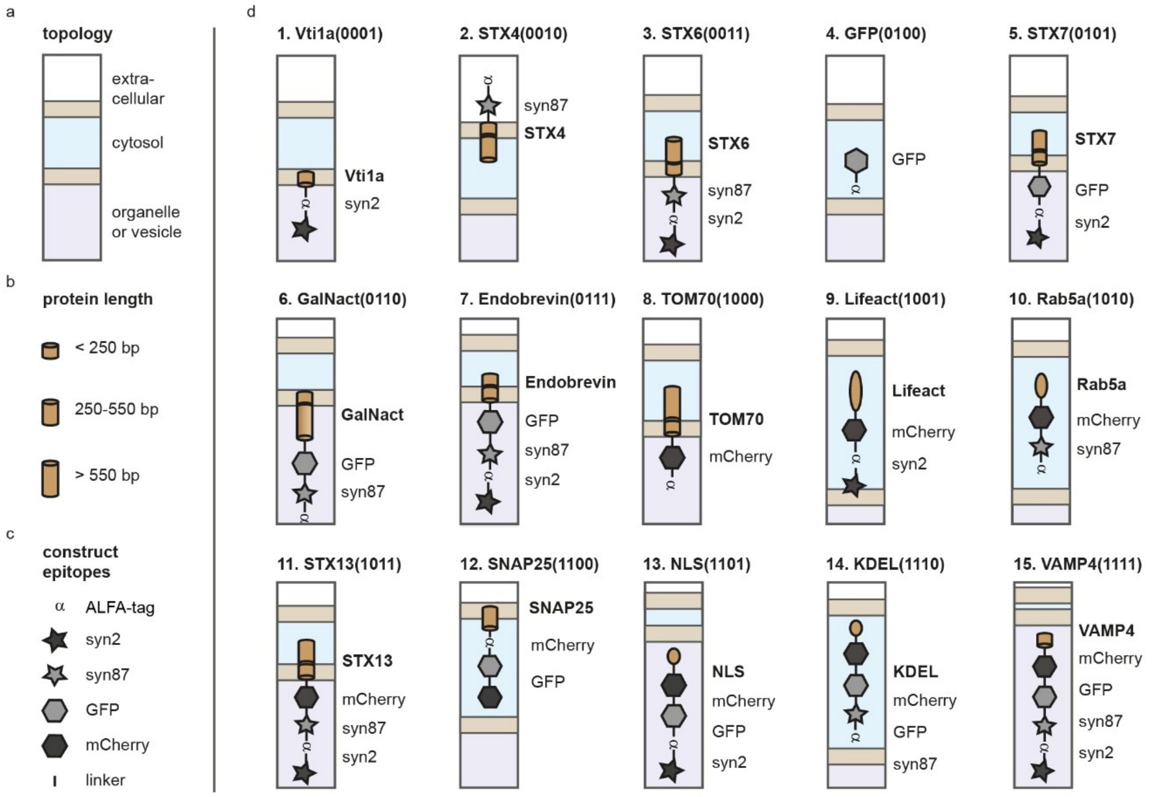


**Extended Data Figure 1: Design and topology of protein constructs.**
**a-c:** Legends for expected topology (a), protein length (b) and construct epitopes (c). **d:** Protein topology schemes for the 15 construct used. Below is a list with detailed information about the respective topology scheme of each construct depicted in b. Uniprot accession numbers (acc.nr.) are available under https://www.uniprot.org/uniprot/.

0. No protein, used for background signals
1. Vti1a(1000): Golgi apparatus and cytosplasmic vesicles Acc.nr.: Q96AJ9
2. STX4(0010): Cytoplasmic, transmembrane and extracellular domain Acc.nr.: Q12846
3. STX6(0011): Cytoplasmic and transmembrane domain Acc.nr.: Q6365
4. GFP(0100): cytoplasmic; the Y66L mutation was added Acc.nr.: C5MKY7
5. STX7(0101): Cytoplasmic, transmembrane and vesicular domain Acc.nr.: O70257
6. GalNacT(0110): Cytoplasmic, transmembrane and luminal domain Acc.nr.: Q10471
7. Endobrevin(0111): Cytoplasmic, transmembrane and vesicular domain Acc.nr.: Q9BV40
8. TOM70(1000): Michondrial, transmembrane and cytoplasmic domain Acc.nr.: P07213
9. Lifeact(1001): Found in cytoplasm and cytoskeleton Acc.nr.: Q08641
10. Rab5a(1010): Found in cell membranes and cytosol Acc.nr.: P20339
11. STX13(1011): Cytoplasmic, transmembrane and vesicular domain Acc.nr.: G3V7P1
12. SNAP25(1100): Cytoplasmic and membrane associated Acc.nr.: P60881
13. NLS(1101): Found in nucleus Acc.nr.: P03070
14. KDEL(1110): Only in combination with ER protein in ER, here in cytosol Acc.nr.: n/a
15. VAMP4(1111): Transmembrane domain Acc.nr.: D4A5

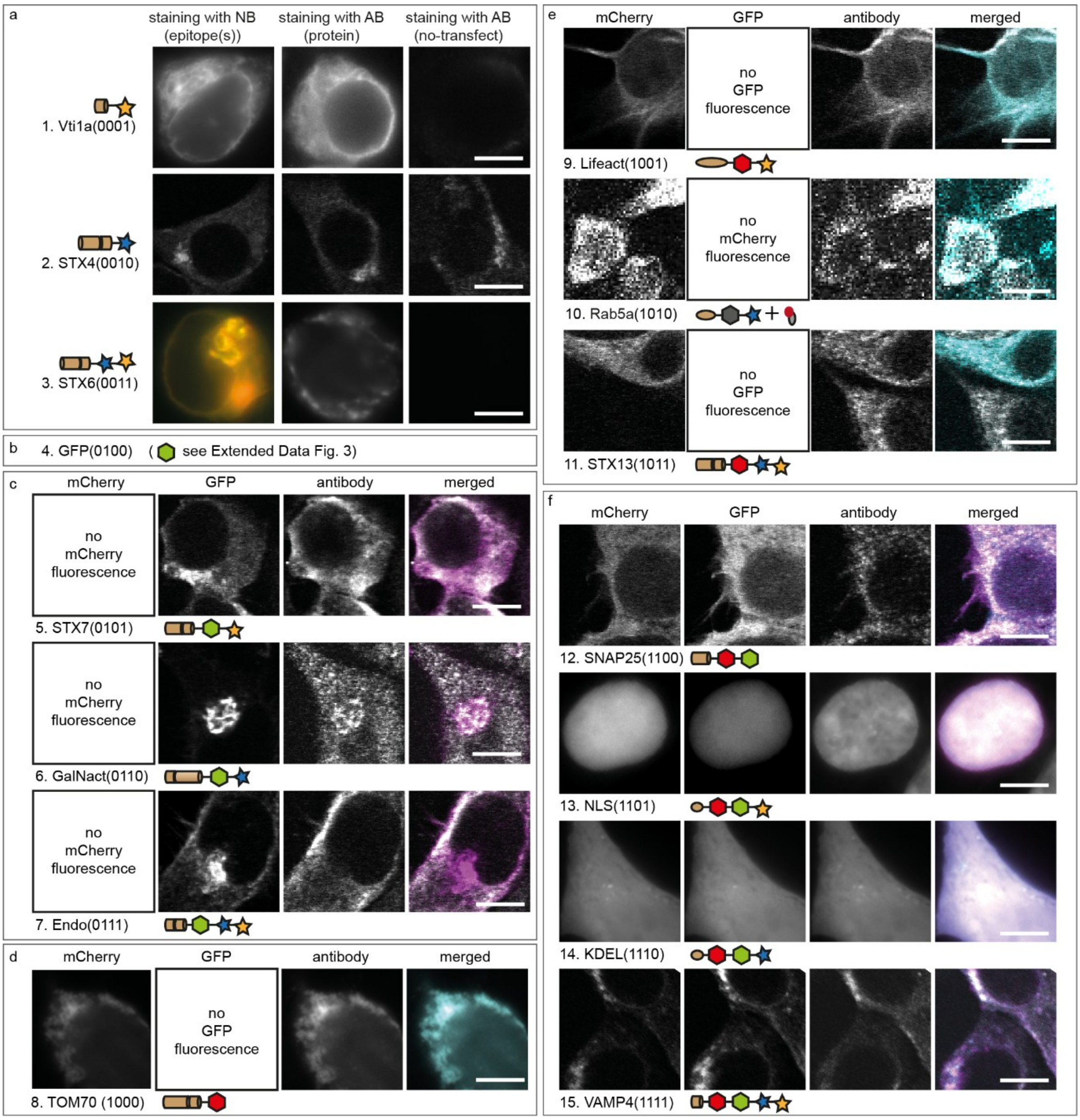


**Extended Data Figure 2: A visualization of nanobarcode-carrying proteins using antibodies.** We validated the correct nanobarcoding and expression of the protein constructs by simultaneous visualization of the nanobarcodes and their respective endogenous epitope counterparts. The nanobarcodes were visualized by imaging their GFP or mCherry fluorescence (relying on constructs lacking the Y/L mutations of the chromophores), or by nanobody stainings, for barcodes lacking GFP or mCherry. We immunostained the respective proteins of interest with antibodies directed against protein-specific epitopes. **a:** All protein constructs lacking mCherry or GFP fluorescence. **b, c:** Protein constructs with GFP fluorescence. These proteins exhibit a strong localization to the perinuclear area, where antibodies penetrate more poorly than nanobodies[1]. See Extended Data Fig. 3 for nanobody staining of the nanobarcode epitopes. **d, e:** All protein constructs having a fluorescent mCherry epitope. **f:** All protein constructs having mCherry and GFP fluorescence. To visualize the target protein component of the nanobarcoded-proteins, Cy5-coupled secondary antibodies were used. Scale bar, 10 µm.

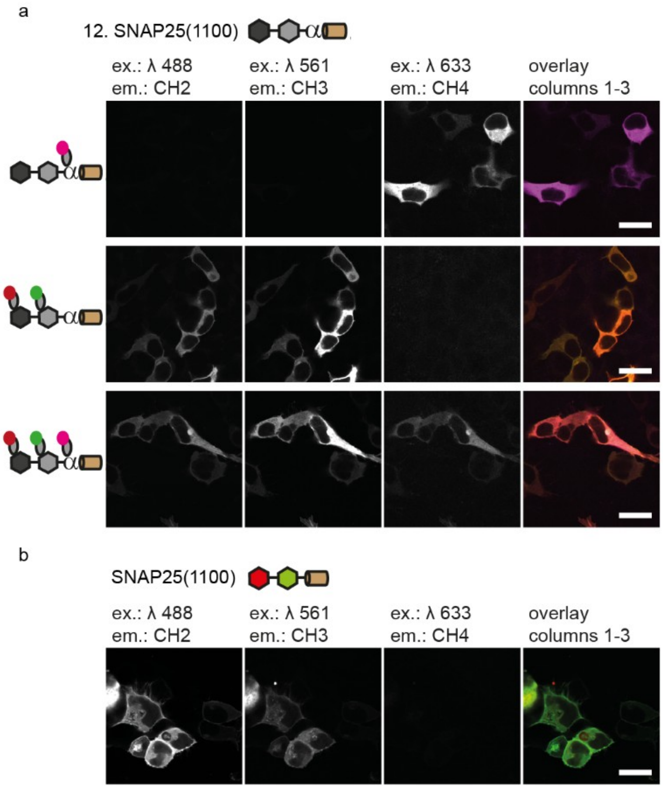
a
12. SNAP25(1100)
ex.: λ 488
em.: CH2
ex.: λ 561
em.: CH3
ex.: λ 633
em.: CH4
overlay
columns 1-3
b
SNAP25(1100)
ex.: λ 488
em.: CH2
ex.: λ 561
em.: CH3
ex.: λ 633
em.: CH4
overlay
columns 1-3

**Extended Data Figure 3: Simultaneous nanobody staining of multiple epitopes. a:** Non-fluorescent epitopes of SNAP25(1100) are recognized successfully by the corresponding nanobodies, independent of the number of nanobodies used. The SNAP25(1100) construct is successfully stained when using the NbALFA only (first row), when using the anti-GFP and anti-RFP nanobodies (second row) or when using the anti-GFP, anti-RFP and anti-ALFA nanobodies (third row). The corresponding nanobarcode epitopes are detected, which indicates that there is no substantial steric hindrance between the nanobodies (which is expected due to the small size of the nanobodies). The morphological features of the cells are similar to cells transfected with a SNAP25 construct containing fluorescent epitopes (b), which indicates that these constructs are comparable and therefore suitable for our investigations, as shown in Fig. 2 and 3. Scale bars, 30 µm.

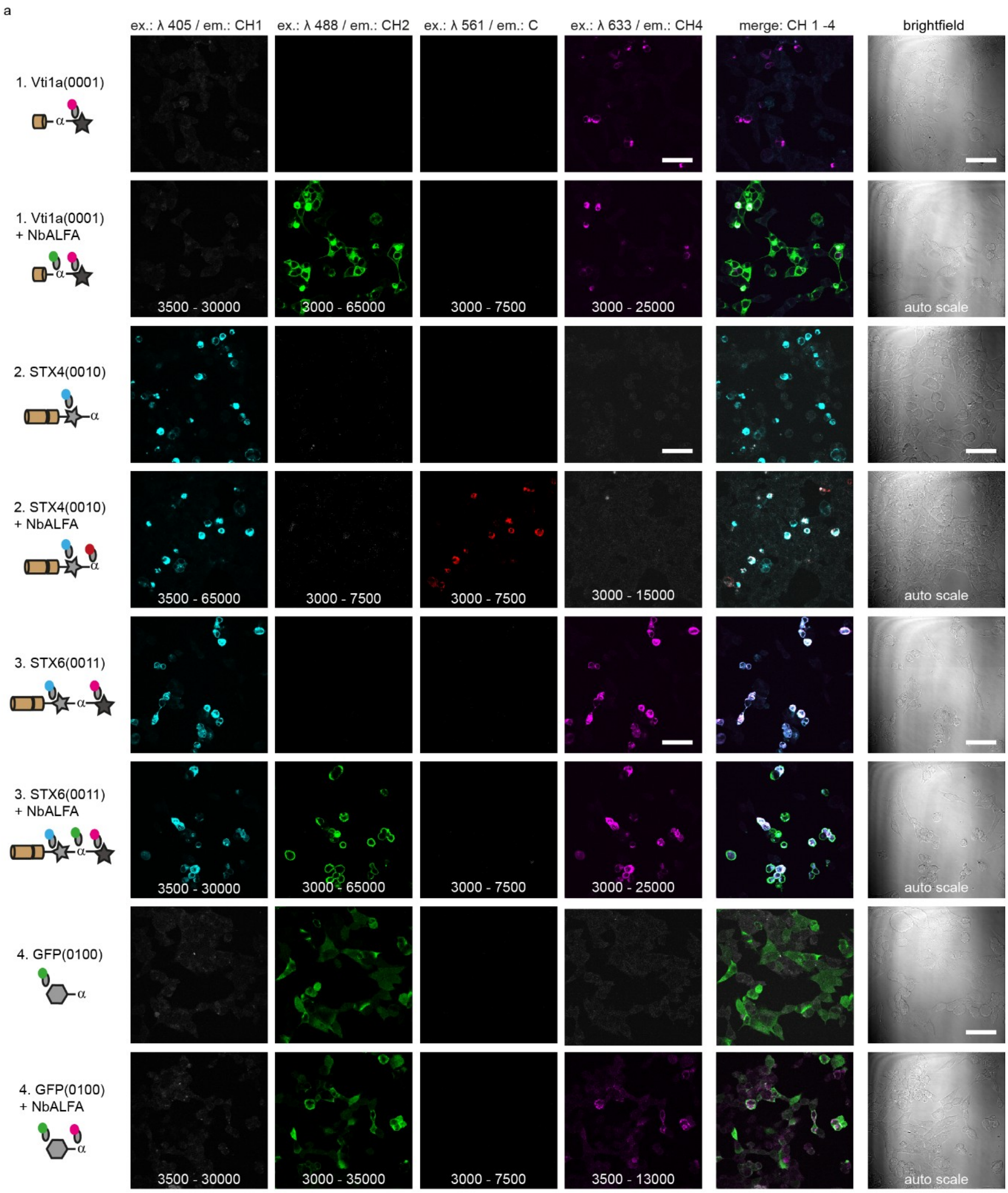
a
ex.: λ 405 / em.: CH1
ex.: λ 488 / em.: CH2
ex.: λ 561 / em.: C
ex.: λ 633 / em.: CH4
merge: CH 1 -4
brightfield
1. Vti1a(0001)
1. Vti1a(0001) + NbALFA
2. STX4(0010)
2. STX4(0010) + NbALFA
3. STX6(0011)
3. STX6(0011) + NbALFA
4. GFP(0100)
4. GFP(0100) + NbALFA
3500 - 30000
3000 - 65000
3000 - 7500
3000 - 25000
auto scale
3500 - 65000
3000 - 7500
3000 - 7500
3000 - 15000
auto scale
3500 - 30000
3000 - 65000
3000 - 7500
3000 - 25000
auto scale
3500 - 30000
3000 - 35000
3000 - 7500
3500 - 13000
auto scale

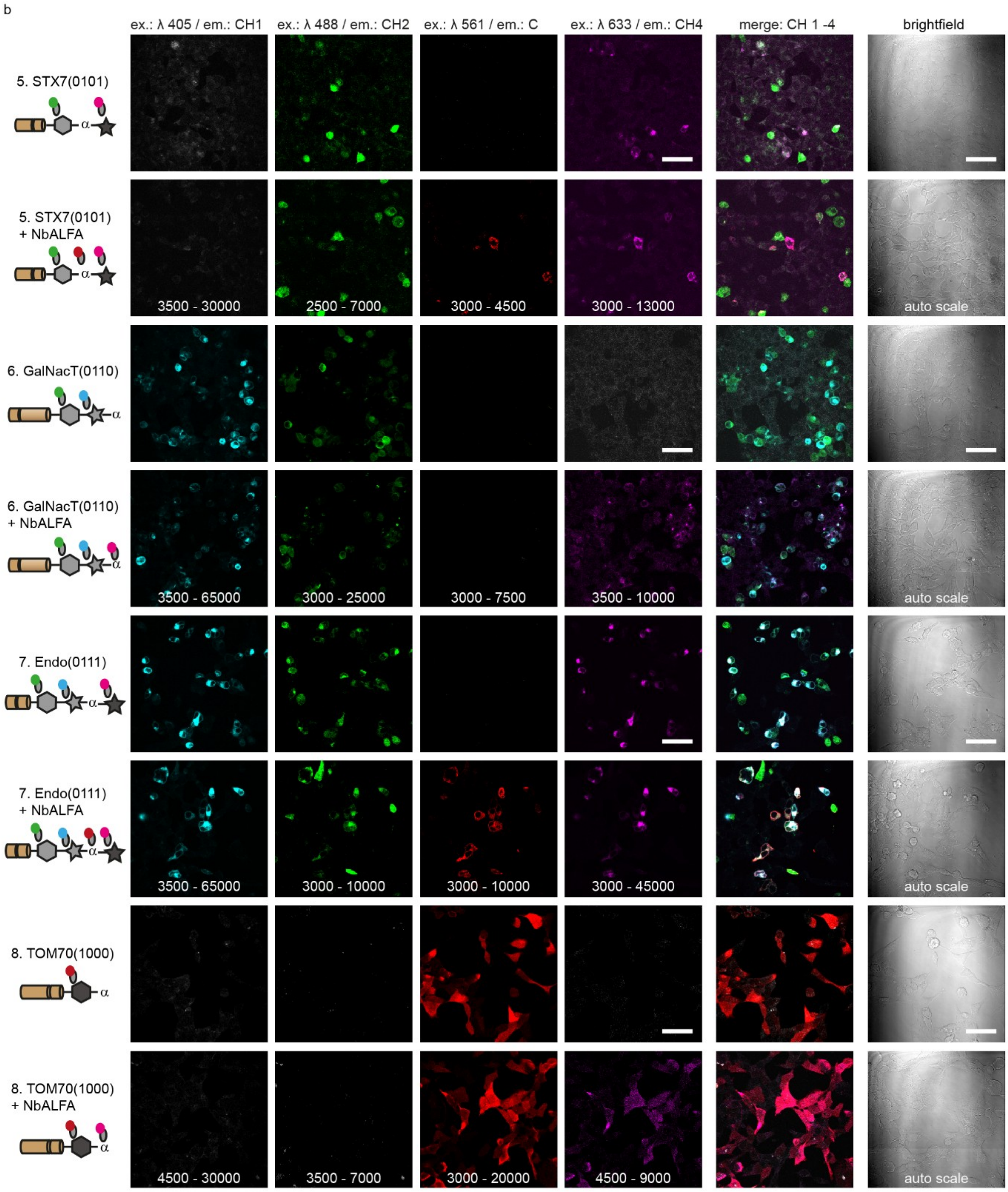
b
ex.: λ 405 / em.: CH1
ex.: λ 488 / em.: CH2
ex.: λ 561 / em.: C
ex.: λ 633 / em.: CH4
merge: CH 1 -4
brightfield
5. STX7(0101)
5. STX7(0101) + NbALFA
3500 - 30000
2500 - 7000
3000 - 4500
3000 - 13000
auto scale
6. GalNacT(0110)
6. GalNacT(0110) + NbALFA
3500 - 65000
3000 - 25000
3000 - 7500
3500 - 10000
auto scale
7. Endo(0111)
7. Endo(0111) + NbALFA
3500 - 65000
3000 - 10000
3000 - 10000
3000 - 45000
auto scale
8. TOM70(1000)
8. TOM70(1000) + NbALFA
4500 - 30000
3500 - 7000
3000 - 20000
4500 - 9000
auto scale

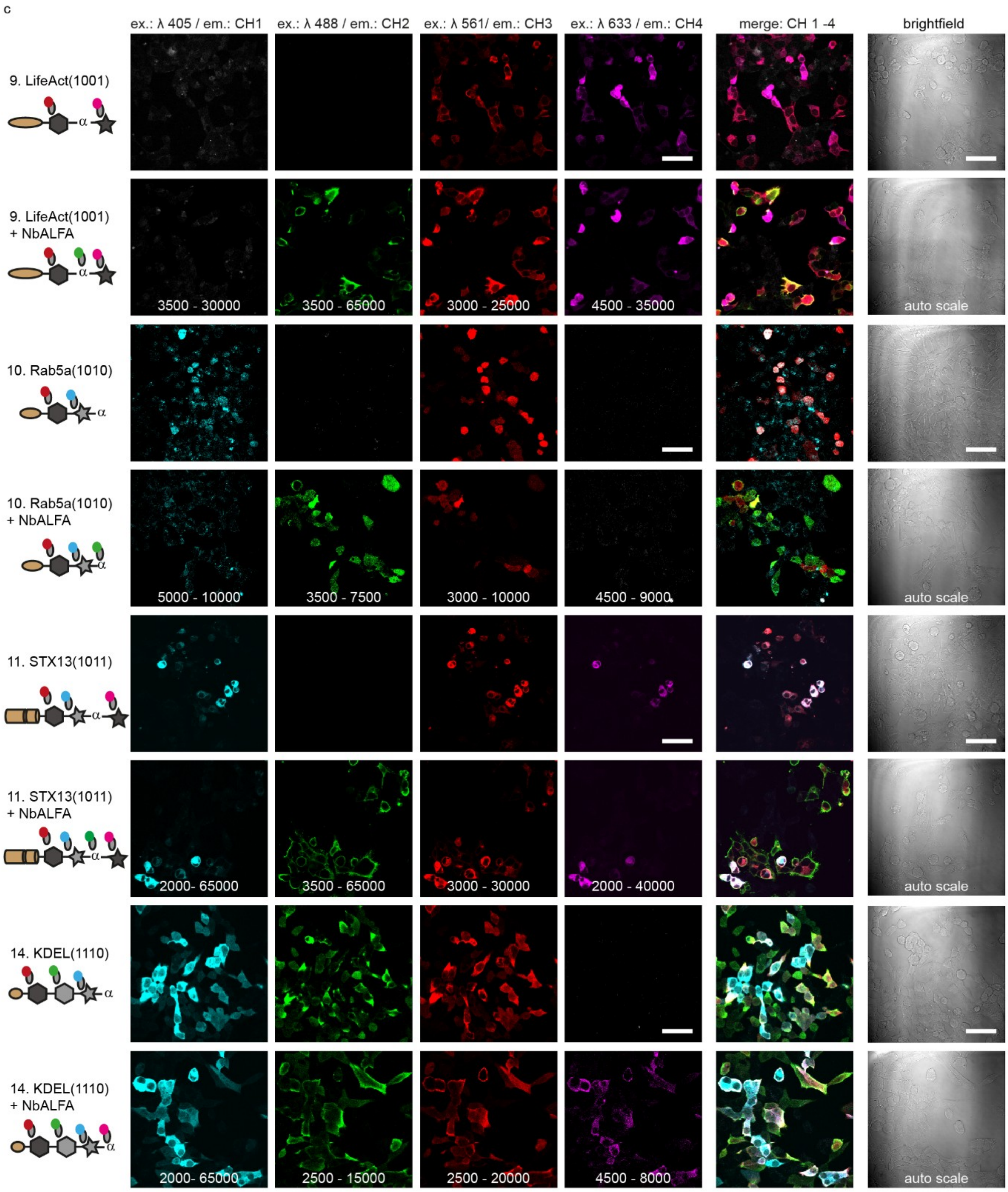
c
ex.: λ 405 / em.: CH1
ex.: λ 488 / em.: CH2
ex.: λ 561/ em.: CH3
ex.: λ 633 / em.: CH4
merge: CH 1 -4
brightfield
9. LifeAct(1001)
9. LifeAct(1001) + NbALFA
3500 - 30000
3500 - 65000
3000 - 25000
4500 - 35000
auto scale
10. Rab5a(1010)
10. Rab5a(1010) + NbALFA
5000 - 10000
3500 - 7500
3000 - 10000
4500 - 9000
auto scale
11. STX13(1011)
11. STX13(1011) + NbALFA
2000- 65000
3500 - 65000
3000 - 30000
2000 - 40000
auto scale
14. KDEL(1110)
14. KDEL(1110) + NbALFA
2000- 65000
2500 - 15000
2500 - 20000
4500 - 8000
auto scale

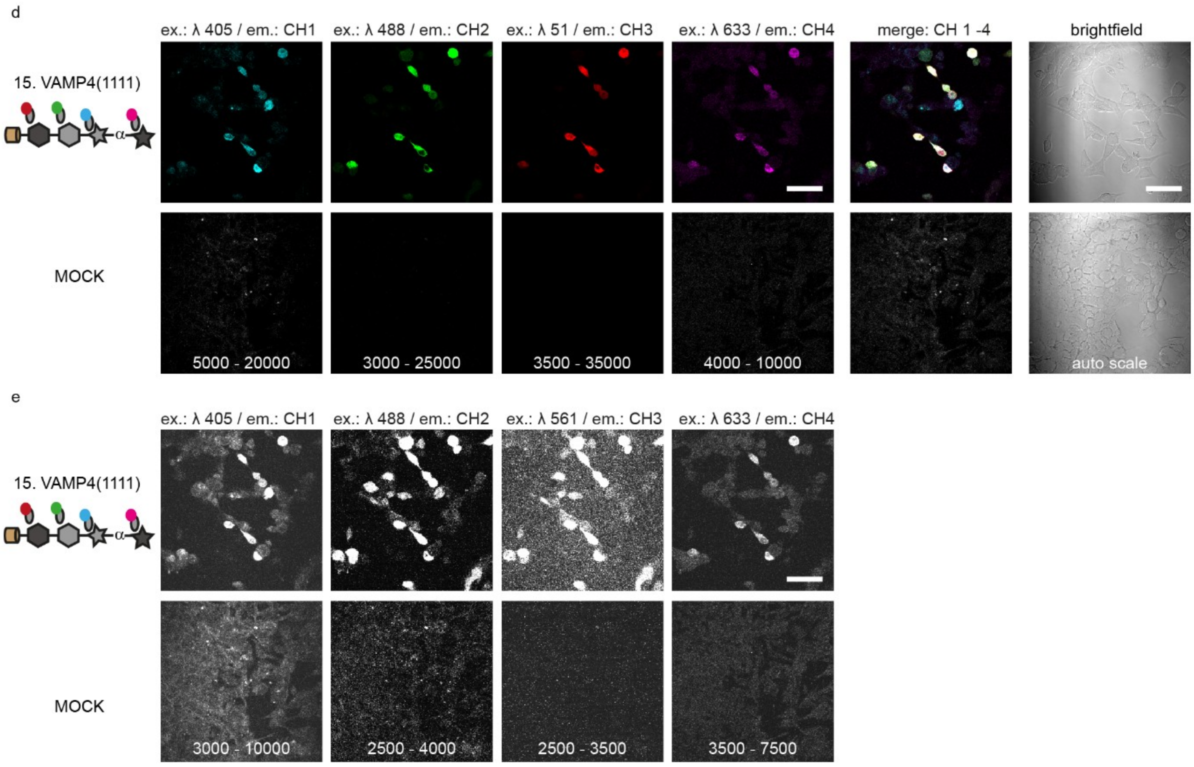


**Extended Data Figure 4: Visualization of fifteen nanobarcode epitopes using four spectrally distinct nanobodies. a-d:** Nanobody-based identification of the four genetically-encoded nanobarcode-epitopes mCherry(Y71L), GFP(Y66L), syn87 and syn2 and the ALFA-tag epitope by their corresponding nanobodies NbRFP, NbEGFP, NbSyn87, NbSyn2 and NbALFA. Scaling was optimized for each protein. **d:** VAMP4(1111) example (all epitopes present) and negative control condition: mock transfection (no DNA, no epitopes present) using same intensity scale. **e:** as in d, now with upscale intensities. Scale bar 50 µm.

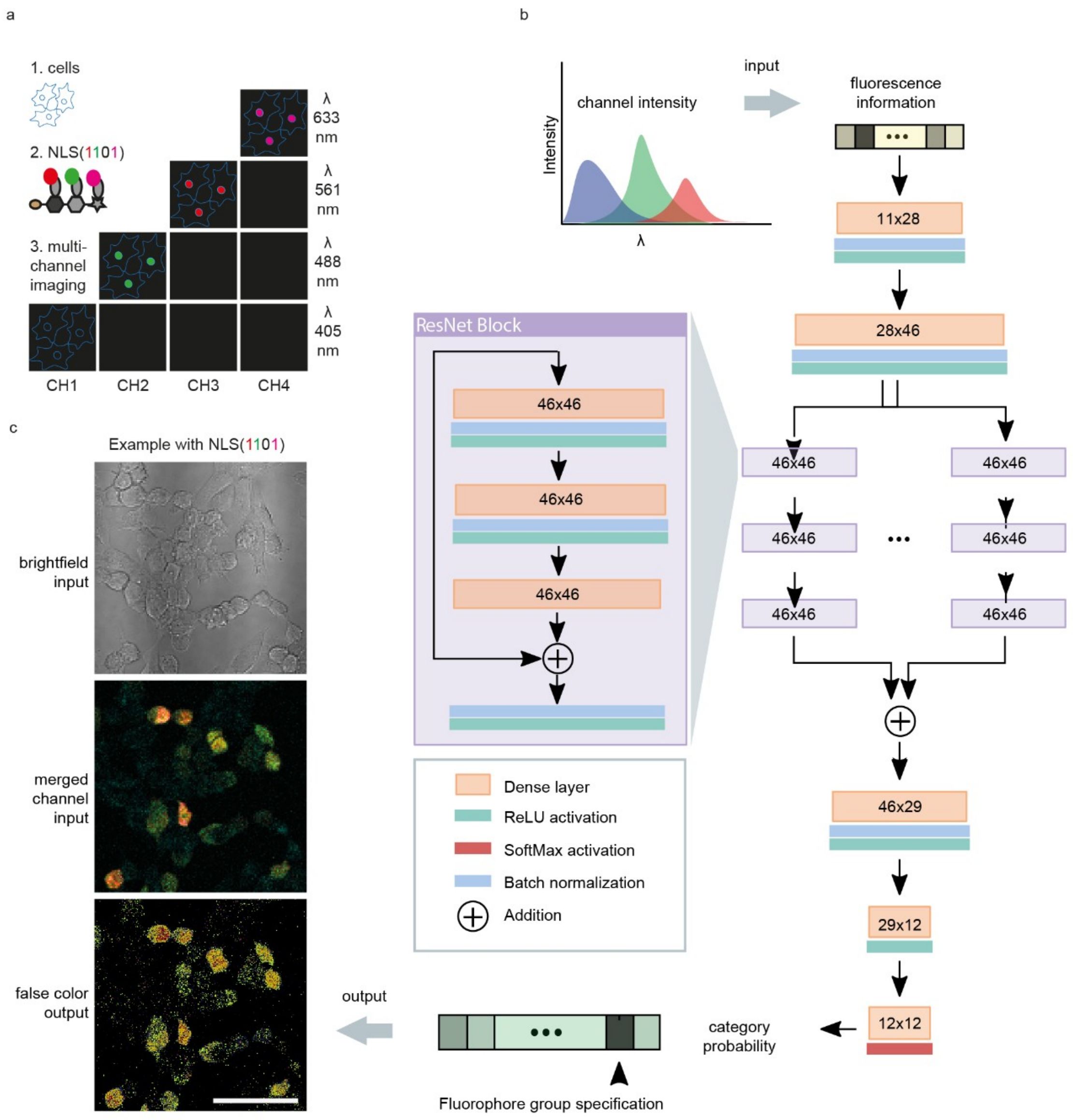


**Extended Data Figure 5: Principles of neural-network-based identification of nanobarcode-proteins. a:** Schematic representation of experimental protocol for obtaining multi-channel images of HEK293 cells transfected with a single protein construct. HEK293 cells are seeded (1) and transfected with the necessary DNA plasmids. After an incubation of at least 14 hours, the HEK293 cells, now expressing the protein constructs, are fixed and stained with nanobodies (2). Multi-channel images from the respective cells (3) are used for the training of a neuronal network. Wavelengths of excitation lasers used: λ = 405 nm, λ = 488 nm, λ = 561 nm and λ = 633 nm. Emission channels used: 417-485 nm (CH1), 495-553 nm (CH2), 573-631 nm (CH3) and 641-729 nm (CH4). **b:** Architecture of the deep neural network used for protein identification from channel intensity values pertaining to each pixel. For the dense layers, given numbers indicate input and output dimensions. The network contains four parallel branches in the middle (two are shown), the outputs of which are summed and processed by the final layers. The branches are composed of sequential ResNet blocks with skip connections bypassing triplets of layers, as shown in the expansion panel to the left (further details in Methods section “Deep neural network-based protein identification”). **C:** The output probability distributions of the network are used to render false-color images that contain information on the identified proteins in each pixel. Scale bar: 50 µm.

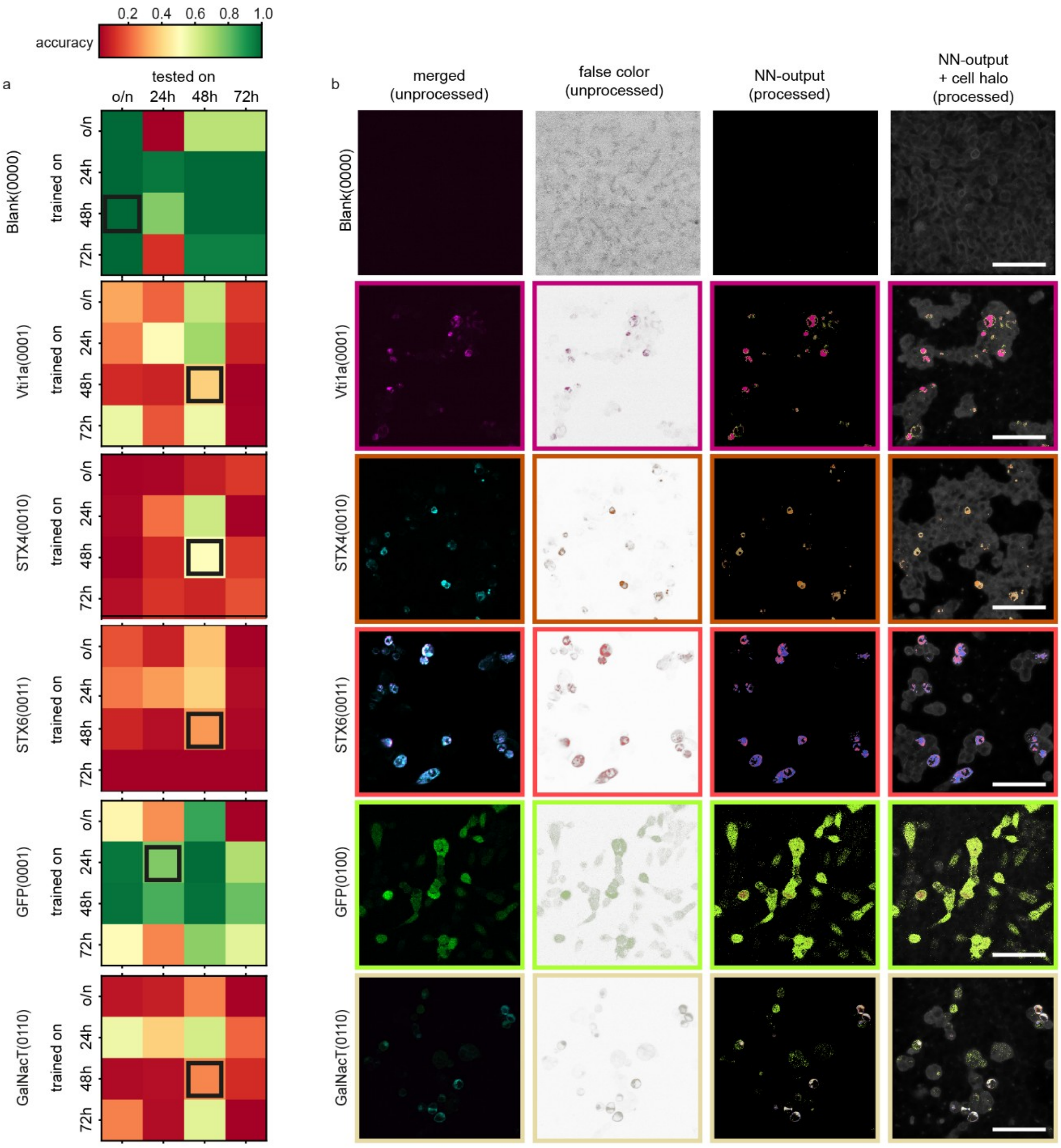

accuracy
0.2 0.4 0.6 0.8 1.0
a
tested on
o/n 24h 48h 72h
trained on
Blank(0000)
Vti1a(0001)
STX4(0010)
STX6(0011)
GFP(0001)
GalNacT(0110)
b
merged (unprocessed)
false color (unprocessed)
NN-output (processed)
NN-output + cell halo (processed)
Blank(0000)
Vti1a(0001)
STX4(0010)
STX6(0011)
GFP(0100)
GalNacT(0110)

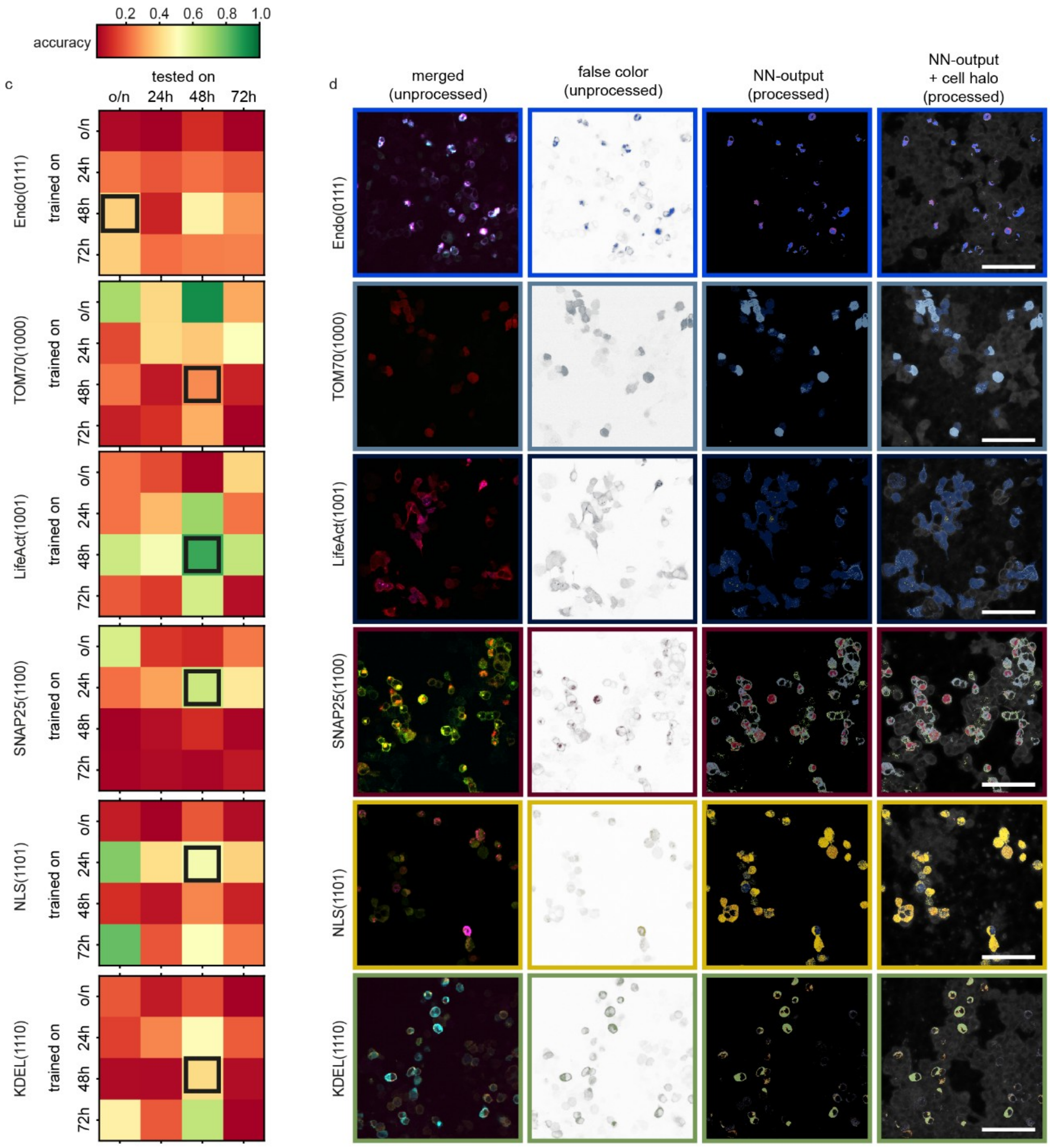


**Extended Data Figure 6: Neural-network analysis results. a**: The prediction accuracy matrix of trained neural networks, estimated over all the images in the dataset. To increase the complexity of the training and testing procedure, we expressed each construct for different time periods, and we then trained and tested the neural networks with all of these different datasets. Each row corresponds to a separate network that has been trained solely on the given dataset. Columns are the average pixel-wise prediction accuracy, assuming that all the pixels picked by the network in an image should belong to the protein with which the cells have been transfected. The given accuracy values may include effects of mis-expressed proteins, weak fluorescence signals and imaging noise. **b**: From left to right, first column: merged channels (405 nm/CH1, 488 nm/CH2, 561 nm/CH3, 633 nm/CH4), before being processed by the network. Second column: images produced by assigning false colors to bright pixels, assuming that all the proteins in the image exactly match the given nanobarcode. Third column: output of the neural network, with each pixel given the false color representing the protein picked by the network. Colors are scaled based on class probabilities (Fig. 2). Fourth column: false color output of the network overlaid on the gray "cell halos" produced from the brightfield images. Brightfield images have been processed to remove noise and background gradients and to enhance the contrast. **c, d**: as a and b, for additional nanobarcode proteins.

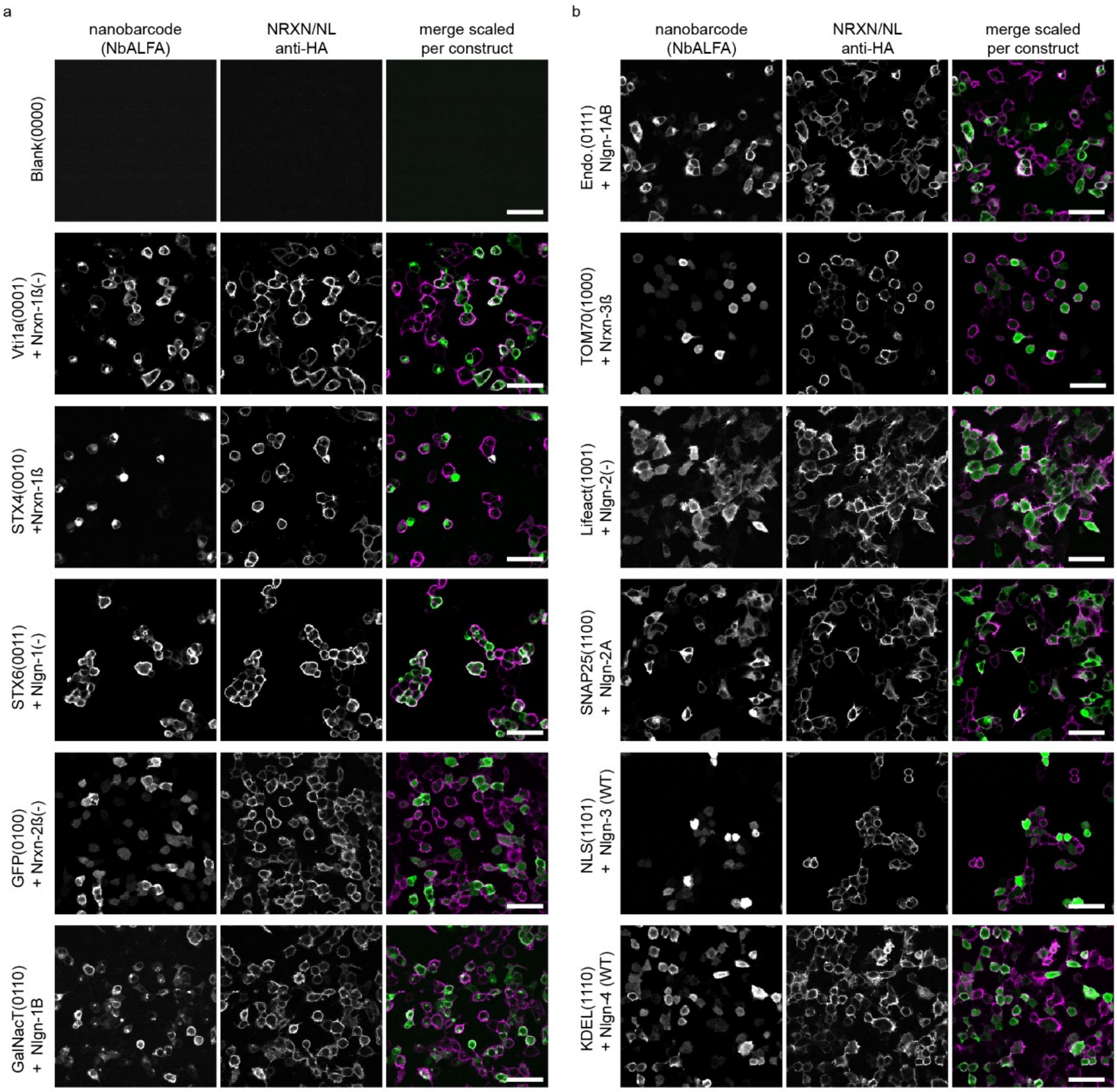


**Extended Data Figure 7: Validation of transfection and expression of protein constructs within HEK293 cells.** To enable an analysis of Nrxn and Nlgn pairing, we co-expressed different Nrxn and Nlgn constructs with specific nanobarcode proteins. This enables us to provide the different Nrxn- and Nlgn-containing cells with a recognizable identity, without having to modify the additional proteins themselves by nanobarcode tagging. However, this implies that we need to verify whether the majority of Nrxn- or Nlgn-expressing cells also express the respective nanobarcode proteins. **a, b:** Nanobody staining with anti-ALFA-Atto488 reveals cells expressing protein constructs with nanobarcodes. Antibody staining with mouse-anti-HA and Cy3-anti-mouse reveals cells expressing NRXN or NL constructs with HA-tags. An overlay of both signals (anti-ALFA in green and anti-HA in magenta) indicates double-transfected cells, which make up the majority of all cells. Scale bar: 50 µm.

| | Antibody | Target protein | Species | Dilution from stock | Company |
|---|---|---|---|---|---|
| 01 | Anti-Vti1a | Vti1a | Rabbit polyclonal | ICC 1:500 | Synaptic System |
| 02 | Anti-Syntaxin 4 | Syntaxin 4 | Rabbit polyclonal | ICC 1:300 | Synaptic Systems |
| 03 | Anti-Syntaxin 6 | Syntaxin 6 | Mouse polyclonal | ICC 1:500 | Synaptic Systems |
| 04 | GFP, does not have a target protein, see Extended Data Figure 3 for validation of epitope | | | | |
| 05 | Anti-Syntaxin 7 | Syntaxin 7 | Rabbit polyclonal | ICC 1:300 | Synaptic Systems |
| 06 | Anti-GM130 | GM130 | Rabbit polyclonal | ICC 1:300 | Sigma-Aldrich |
| 07 | Anti-Endobrevin | Endobrevin | Rabbit polyclonal | ICC 1:300 | Synaptic Systems |
| 08 | Anti-TOMM20 | Sigma | Mouse monoclonal | ICC 1:200 | Sigma-Aldrich |
| 09 | Anti-Beta-Actin | Beta-Actin | Mouse polyclonal | ICC 1:75 | Sigma-Aldrich |
| 10 | Anti-Rab5a | Rab5a | Rabbit monoclonal | ICC 1:250 | Abcam |
| 11 | Anti-Syntaxin 12/13 | Syntaxin 12/13 | Rabbit polyclonal | ICC 1:300 | Synaptic System |
| 12 | Anti-SNAP 25 | SNAP 25 | Rabbit polyclonal | ICC 1:500 | Synaptic Systems |
| 13 | Anti-NFkappaB p65 | NLS | Rabbit monoclonal | ICC 1:200 | Biomol / Rockland |
| 14 | Anti-KDEL | KDEL | Mouse monoclonal | ICC 1:500 | Enzo Life Sciences |
| 15 | Anti-VAMP 4 | VAMP 4 | Rabbit polyclonal | ICC 1:300 | Synaptic Systems |
| 16 | Anti-Rabbit IgG Cy5 | Anti-Rabbit IgG | Donkey | ICC1:500 | Dianova |
| 17 | Anti-Mouse IgG Cy5 | Anti-Mouse IgG | Donkey | ICC 1:500 | Dianova |

**Extended Data Table 1: Information about antibodies used for target protein validation purposes.** The first fifteen antibodies listed are primary antibodies. Antibodies sixteen and seventeen are secondary antibodies. See Methods section for further information about the staining procedures.
